# Interaction of attention and localization of pain maxima shift the balance between lateral inhibition and spatial facilitation in nociceptive processing

**DOI:** 10.64898/2026.03.14.711815

**Authors:** Jakub Nastaj, Tibor M. Szikszay, Jacek Skalski, Kerstin Luedtke, Robert C. Coghill, Wacław M. Adamczyk

**Affiliations:** Institute of Physiotherapy and Health Sciences, Academy of Physical Education, Katowice, Poland; University of Luebeck, Institute of Health Sciences, Department of Physiotherapy, Pain and Exercise Research Luebeck (P.E.R.L.), Luebeck, Germany; Pediatric Pain Research Center (PPRC), Cincinnati Children’s Hospital Medical Center, Cincinnati, Ohio, USA; Division of Behavioral Medicine and Clinical Psychology, Cincinnati Children’s Hospital Medical Center, Cincinnati, Ohio, USA; Department of Pediatrics, University of Cincinnati College of Medicine, Cincinnati, Ohio, USA

**Keywords:** lateral inhibition, spatial summation of pain, nociception, attention, diffuse noxious inhibitory control

## Abstract

Spatial interactions among noxious inputs can enhance or suppress pain, but the conditions determining whether spatially organized nociceptive processing is dominated by facilitation or inhibition remain unclear. The present study examined whether attentional tuning and localization of pain maxima regulate the balance between lateral inhibition (LI) and spatial summation of pain (SSP). Thirty healthy participants completed a within-participant psychophysical experiment with electrocutaneous stimulation across LI, sham-control, diffuse noxious inhibitory control, and SSP trials. The spatial configuration of noxious input was manipulated by varying the number and arrangement of activated electrodes. Depending on the attentional instruction, participants continuously rated either pain intensity attributed to a target electrode or overall pain from all activated electrodes using an electronic visual analogue scale. Secondary measures assessed target-pain extraction, attentional focus, and localization of the pain maximum. Rather than observing the predicted inhibitory profile, pain ratings increased suggesting a predominance of facilitatory spatial interactions that were strongly modulated by attention and the perceived location of the pain maximum. When the pain maximum was localized outside the target electrode, facilitation was absent and overall pain intensity was lower. Bilateral stimulation of homologous sites elicited SSP, indicating integration of noxious input across body sides. These findings suggest that LI and spatial facilitation are not fixed consequences of stimulus geometry. Instead, their expression depends on the interaction between top-down attentional selection and spatial attribution of the most intense component of pain.

**Summary:** Pain depends on the interaction between the spatial position of its maximum and attentional focus, determining individual facilitatory or inhibitory profiles.

## 1. INTRODUCTION

Across the human neuroaxis, sensory systems employ neurophysiological mechanisms to encode both the intensity and spatial structure of external stimuli. In vision, stimulation of a given neuron elicits excitation of that neuron while producing inhibition in adjacent neurons. This mechanism, known as lateral inhibition (LI), has been extensively studied and, owing to the ON/OFF organization of retina cells, has been linked to contrast enhancement phenomena such as Mach band effects [9,18,30,36,48]. Through such spatial modulation, LI is thought to increase the precision with which the nervous system represents the external environment [16,26,47,59], thereby optimizing interactions within it [8–10].

Converging evidence across sensory modalities suggests that LI represents a general computational principle [35,37,47,67] and may therefore also operate within the human nociceptive system underlying pain perception [20,41,58]. Several psychophysical observations have been interpreted in this framework, including paradoxical findings of equal [3] or reduced [58] pain intensity following stimulation of larger skin areas – effects that partially diverge from classical observations derived from studies on spatial summation of pain (SSP) [13,15,25,53]. At the same time, SSP has been shown to be nonlinear [3,5] and subadditive [1,3,29,40], implying that excitatory nociceptive responses are subject to inhibitory modulation [34,62]. Such inhibition may arise from local lateral interactions or from global mechanisms such as diffuse noxious inhibitory control (DNIC), in which a remote heterotopic noxious stimulus suppresses pain elsewhere [34,54]. Despite these observations, the magnitude and functional relevance of LI within the nociceptive system remain poorly explored, and it remains unclear whether inhibitory spatial effects observed psychophysically reflect genuine lateral interactions or alternative descending control mechanisms such as DNIC.

Studying LI in humans poses substantial methodological challenges. Prior work has largely relied on forced-choice paradigms, such as nociceptive two-point discrimination (2PD), in which increased spatial separation required to discriminate two nociceptive foci is attributed to lateral inhibitory interactions [20,58]. While highly informative, such approaches primarily assess spatial discrimination and may therefore capture only one manifestation of LI in pain perception. Moreover, performance on these tasks is strongly influenced by attentional and decisional processes. Here, we introduce a complementary psychophysical framework designed to examine whether LI modulates pain intensity under attentional conditions. Participants selectively rated pain arising from a single target electrode while adjacent electrodes were concurrently activated in close spatial proximity, enabling controlled spatial extraction of pain. Based on the proposed interactions between spatially adjacent nociceptive inputs, we hypothesized that pain elicited by the target electrode would be reduced when adjacent electrodes were also co-activated, compared with stimulation of the target electrode alone or with sham stimulation. We further hypothesized that this modulatory effect would depend on the number of co-activated (adjacent) electrodes. To dissociate LI from other spatial and nonspecific modulatory processes –including SSP, DNIC, and attentional effects– the study design incorporated dedicated control conditions tailored to isolate lateral inhibitory contributions.

## 2. MATERIAL AND METHODS

### 2.1. General information

The study protocol was approved by the Bioethics Committee at the Academy of Physical Education in Katowice (no. 3-VII/2024) and registered at the Open Science Framework platform using the AsPredicted.org template (https://osf.io/d52rw/). The study was conducted in the certified (ISO) Laboratory of Pain Research. The experiment followed the recommendations of the Declaration of Helsinki [71]. Participants were given written and oral information of the study procedures before informed consent was obtained. Moreover, participants were informed that they could withdraw from the study at any timepoint without any reason and consequences. Each volunteer was offered financial remuneration for their participation (90 PLN).

### 2.2. Study population and eligibility

A group of 33 healthy participants (16 females) from 20 to 38 years (age: mean of 23.60 ± SD 4.30) was recruited (between August and December 2025). The following eligibility criteria were applied: participants had to be pain-free on the day of the study, healthy (self-report), not experienced prolonged pain in the last 24 hours, no chronic pain, no pregnancy, no cardiovascular or neurological disease, no chronic medication use, no mental illness or any systemic disease, no electronic devices in or on the body or unremovable metal objects in area of the both hands, as well as no skin allergies, skin lesions, tattoos or reported sensory abnormalities in close proximity to the both hands.

### 2.3. Outcome measures

#### 2.3.1. Primary outcome

The main outcome measure in this experiment was the real time pain intensity rating – participants rated the perceived pain intensity during the nociceptive stimulation by using an electronic Visual Analogue Scale (eVAS) which was displayed on the computer’s screen. The scale was ranging from “0” (no pain sensation) to “100” (worst pain sensation imaginable) [4,52].

#### 2.3.2. Secondary outcomes

After termination of each nociceptive stimulation participants were asked to rate their own ability to extract pain sensations in accordance with the instructed rating task. These ratings were collected by using the eVAS, ranging from “0” (impossible) to “100” (entirely possible). The aim of this measure was to assess the extent to which participants were able to extract the pain sensation arising from the target electrode when other electrodes delivering painful sensations were concurrently activated. Following pain extraction rating, participants were asked to rate their ability to focus attention on pain sensations, depending on the instructed rating task. These ratings were also collected by using the eVAS, ranging from “0” (impossible) to “100” (entirely possible). Specifically, this measure evaluated participants’ ability to focus their attention on pain originating from the target electrode in the presence of nociceptive input from other electrodes. After each trial participants also indicated under which electrodes pain was the most intense, which was used as a proxy of perceived locus of pain maximum.

#### 2.3.3. Additional variables

Further variables were collected to characterize the study sample. These included: age, sex (assigned at birth), handedness, body height (cm), and body mass (kg), fear of pain as a state – assessed on a “0” (not fear at all) to “10” (highest possible fear) on a Numerical Rating Scale (NRS), and as a trait – measured via the Fear of Pain Questionnaire III (FPQ-III) [42]. Each study visit finished with short survey including three control questions: (i) whether the location of the most intense pain changed during stimulation? (possible answers: YES/NO) (ii) whether the location of the most intense pain during stimulation was felt at the site of the electrodes or between the electrodes (possible answers: YES/NO), and (iii) what was their guess regarding the aim of the experiment – open question.

### 2.4. Apparatus, stimulation, and experimental setup

During the experiment, participants sat upright in front of the computer desk. In total, seven planar-concentric gold-covered (∅ 8 mm) electrodes (WASP, Brainbox Ltd., Cardiff, United Kingdom), separated by 3 cm were placed on the dominant (3 electrodes) and non-dominant hand (4 electrodes). The 3-cm inter-electrode distance was selected to balance functional separation with anatomical consistency. Previous studies conducted on the dorsum of the hand indicate that nociceptive stimuli separated by approximately 3 cm can be perceived as distinct loci [38], supporting the use of this spacing to differentiate adjacent nociceptive inputs [39]. At the same time, the distance kept the stimuli sufficiently close to allow interactions between adjacent inputs, including lateral inhibitory mechanisms. This spacing also ensured that all electrodes remained within the same dorsal hand region across participants, avoiding extension beyond the wrist joint line in individuals with smaller hands.

Electrodes were placed on the dorsal surface of both hands [69,70], in the area between the metacarpophalangeal joints and the wrist crease **(Figure 1A)**. On each hand, the electrodes were arranged in a triangular configuration. On the non-dominant hand, a central electrode was positioned at the centre of the triangle. The orientation of the triangle’s apex was aimed to be counterbalanced across participants such that even subjects had orientation with base of the triangle positioned proximally versus distally to control for potential site-specific inhibitory effects.

**Figure 1.**
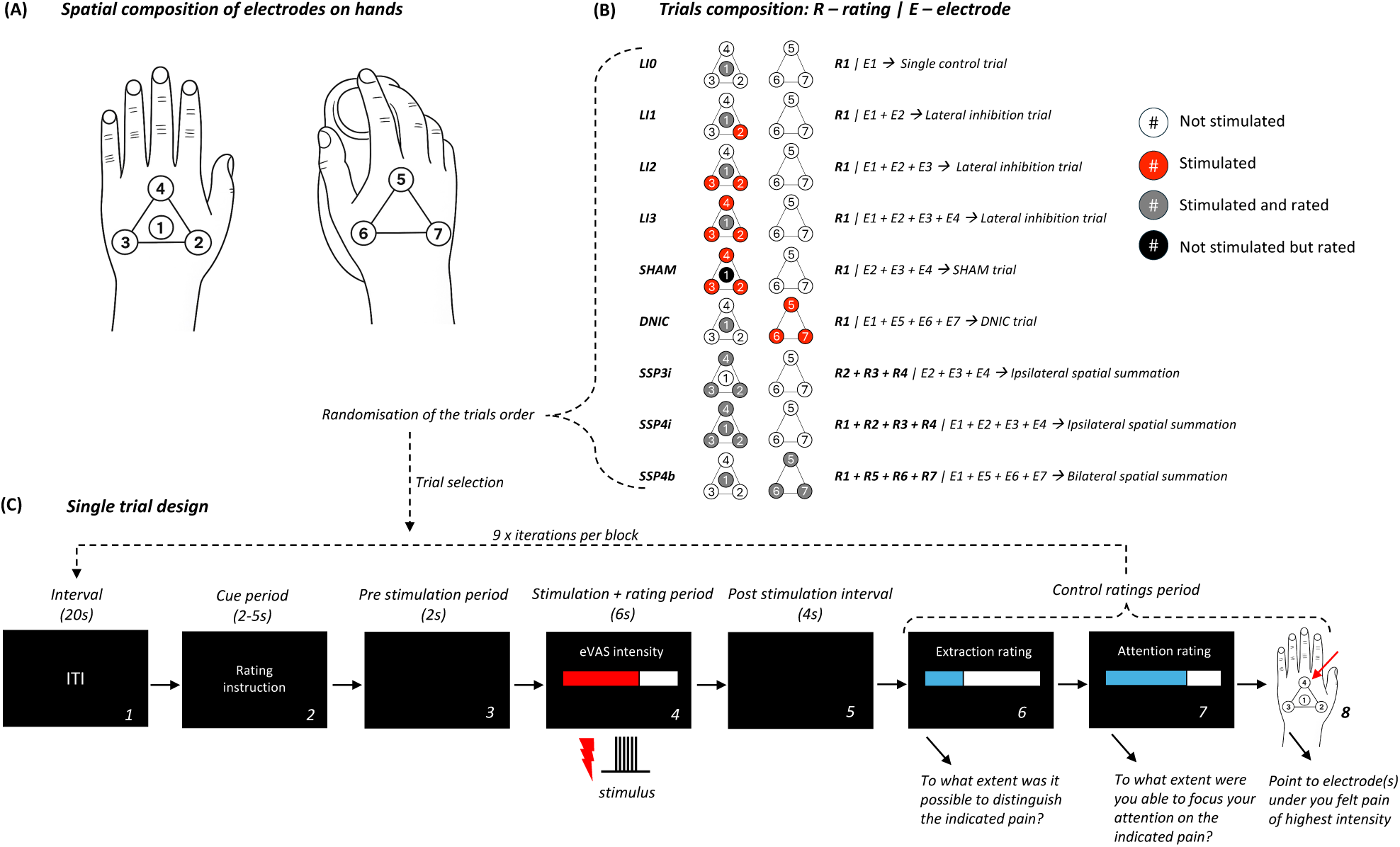
Experimental design. **(A)** Spatial configuration of electrodes. Electrode numbers and their positions on the dorsal side of both hands are indicated in black (circles). **(B)** Types of trials used in the main part of the experiment: The number of electrodes which participants were asked to consider during pain intensity rating are indicated in grey circles. The number of electrodes which were activated but not rated by participants during the trial are indicated in red circles. The electrode which was not activated but rated by participants during the trial is indicated in black circle. The number of electrodes which were not activated during the trial are indicated in white circles. “**R1**” indicates the pain intensity rating from the target middle electrode **E1**. When more electrodes are highlighted under “**R**” it denotes that participants had to provide the overall pain rating from all electrodes activated. The number and identity of the activated electrodes **“E”** (**E1-E7**) are shown in bold along with the corresponding trial name. For example, trial **R1**|**E1** refers to the single control trial (LI0) in which middle electrode **E1** was activated and subjects rated **(R)** pain only from this electrode. **(C)** Single trial design. For each of the two main blocks of trials, trial selection was random: 1) Intertrial interval: 20s between each trial, 2) Task instruction and attentional guidance: A task cue instructed participants to either rate pain from the middle electrode only or to rate overall pain from all active electrodes. For the former, the experimenter pointed to the central electrode location and directed attention to that site, whereas for the latter, participants were asked to evaluate pain globally without focusing on any specific electrode, 3) Two seconds of pre stimulation interval, 4) Stimulation period lasted for six seconds during which pain was continuously rated, 5) four second of post-stimulation interval, 6) Pain extraction rating, 7) Attention rating, 8) Rating of perceived locus of pain maximum: participants indicated electrodes under which they felt most intense pain.

The dominant hand was used to perform the pain intensity rating (as it was involved in stimulation only in two trials), pain extraction rating and attentional rating via a track ball (BIGTrack 2, AbleNet, USA). Trackball was preferred over the mouse to reduce the movement of the hand allowing to provide ratings with mostly single digit. Electronic Visual Analogue Scale and written instructions were displayed on the screen (E900, BENQ, 1280×1024, Taiwan) placed in front of the participant (distance approximately 60 cm). A schematic representation of electrode placement and numbering was provided on the desk in front of the participants for additional reference. The non-dominant hand with the target electrode was covered by an opaque fabric that concealed the position of the electrodes. A dot was marked on the material indicating the position of the target electrode.

To induce pain, electrocutaneous stimuli were delivered via a DS7A constant current stimulator (Digitimer, Welwyn, Garden City, UK) with a maximum voltage capacity of 400 V. Each single stimulus was formed by a series of (rectangular) pulses (200μs duration, 20Hz). Stimulation intensity was fixed at 6 mA across all participants to preserve natural inter-individual variability and avoid calibration-induced distortions of the outcome distribution [6]. Each stimulus formed by pulses lasted 6 seconds. Pulses were distributed to the target electrodes via a D188 remote electrode selector (Digitimer, Welwyn, Garden City, England), which selected the electrode configuration delivering stimulation on a trial-by-trial basis. External digital control of the DS7A and D188 was ensured through a digital/analogue converter device Labjack U3-LV (LabJack Corporation, Lakewood, CO, USA), which was programmed using “u3” Python-3 library. Experimental procedures were operated by the Python 3.12.

### 2.5. Study design

The experiment utilized a within-subject design and involved different experimental and control trials. The trials in the main experimental phase (see section 2.6.4.) differed in relation to the spatial extent of nociceptive input and the type of ratings participants were asked to provide. The spatial extent of nociceptive input was controlled by activation of different numbers of electrodes. The type of ratings consisted of either *directed* pain intensity ratings from a target electrode only, or *overall* pain produced by all electrodes which were activated in a given trial. Participants were asked to continuously rate pain intensity from cued electrode(s) during the entire stimulation period. For simplicity, below, each trial type and its rationale is explained descriptively, and their visual representations are provided on **Figure 1B**.

1. LI trials: in these unilateral trials, participants always provided ratings from a middle target electrode labelled as electrode 1 (E1). These trials involved four different stimulus configurations - either single E1 activation <u>(LI0)</u> or co-activation with adjacent electrode(s): E2 <u>(LI1)</u>, E2 and E3 <u>(LI2)</u>, or E2, E3 and E4 <u>(LI3)</u>. The reasoning behind the choice of LI trials was that adjacent (surrounded) nociceptive input should inhibit the nociceptors activated by a target electrode.
2. SHAM trial: To attempt to isolate LI from nonspecific effects related to expectations and attention, a <u>SHAM</u> trial was included. Subjects rated pain from the middle E1 electrode. However, this electrode was not activated (unknown to participants). Only surrounding electrodes, i.e., E2, E3 and E4 were activated (see **Figure 1A** and **1B**).
3. DNIC trial: To compare LI effects with DNIC, in which remote pain inhibits a target pain, the <u>DNIC</u> trial was utilized. Subjects rated pain from E1, however, stimulation involved E1 and three electrodes positioned on the contralateral hand (E5, E6 and E7).
4. SSP trials: These trials involved overall pain ratings from different distributions of active electrodes in three different configurations. Ipsilateral SSP trials included active electrodes E2, E3 and E4 (<u>SSP3i</u>) or E1, E2, E3 and E4 (<u>SSP4i</u>). Bilateral SSP trials included active electrodes E1, E5, E6 and E7 <u>(SSP4b)</u>.

### 2.6. Experimental procedures

The experiment started from the briefing and consenting procedure. The examiner verified the inclusion and exclusion criteria and prepared participants for the study procedures. The study procedure consisted of five phases: i) procedure instruction, ii) scale training and familiarization to electrocutaneous stimuli, iii) familiarization to experimental trials, iv) main phase of the study (two blocks of nine trials including experimental and control trials, see **Figure 1B** and **1C**), and v) the exit phase.

#### 2.6.1 Preparation and initial briefing

After cleaning the dorsal skin of the hands with an abrasive gel (NuPrep^®^, Weaver and Company, Aurora, CO, USA) and an alcohol solution, dry electrodes were attached to the participants’ hands, followed by a procedural briefing. Participants were instructed that they would be asked to rate pain intensity in real time (i.e. during the stimulation periods), either from a specific location corresponding to the site of the target electrode or as the overall pain experienced. Such procedure/ratings were successfully used previously, however, in relation to stimuli which were either separated by a larger distance (10 cm) [56] or when the stimulus formed a contiguous area [2]. The use of the eVAS for pain intensity ratings was explained, including instructions (see **Appendix S5**) on pain extraction and attentional focus on the pain experience. Each of the three tasks was presented to allow participants to familiarize themselves with providing ratings using the eVAS and operating the trackball. Following the briefing, and after participants confirmed their understanding of the tasks and rating procedures, the subsequent phase of the experiment commenced.

#### 2.6.2 Scale training and familiarization to electrocutaneous stimuli (block 1)

To familiarize participants with nociceptive stimulation and to allow training in real-time pain intensity ratings, participants completed four trials in which only a single (yet different) electrode was activated. During this phase, stimulation was delivered exclusively via electrodes positioned on the nondominant hand (E1, E2, E3 and E4). The order of electrode activation was fully random. After each trial, participants were asked to report which electrode produced the pain, to assess their ability to accurately localize the site of stimulation on the hand dorsum. This phase also allowed to verify if there were any differences in skin sensitivity between the stimulation sites.

#### 2.6.3 Familiarization to experimental trials (block 2)

To familiarize participants with rating pain intensity from the site of the target electrode as well as with the overall pain experience during experimental trials, participants completed four clearly distinct trials (LI3, SHAM, DNIC, and SSP4b). The order of trial presentation was randomized. After each trial, participants reported whether the intensity of stimulation from any electrode was perceived as more intense than that from the others.

#### 2.6.4. Main experimental phase (blocks 3 and 4)

The main experimental phase was divided into two stimulation blocks. In each block, participants received nine stimulation trials, one trial per experimental condition (**Figure 1B and 1C**) such that eventually each trial type was repeated twice. The order of trials within each block was fully random. Overall, participants completed a total of 18 stimulation trials during the main experimental phase. The inter-trial interval was 20 s, and a 2-min break was provided between the two stimulation blocks.

#### 2.6.5. Single trial design

Each trial consisted of the following phases (**Figure 1C: 1-8)**: 1) Inter-trial interval: Between each trial a 20-seconds interval was used, 2) Task instruction and attentional guidance: A task cue was displayed on the screen instructing participants to either “Rate pain from the middle electrode only” or “Rate overall pain from all electrodes.” When the instruction required rating pain intensity originating from the site of the middle target electrode, the examiner approached the participant’s hand, pointed the location of the central electrode (marked with a dot on fabric that covers hand), and instructed the participant to focus on sensations arising from that specific site. When the instruction required rating overall pain, participants were reminded to evaluate the pain experience globally, without focusing on any specific electrode, 3) Pre stimulation period: An eVAS for pain intensity was presented on the screen, and rating acquisition began. The scale remained visible for 12 seconds, 4) Stimulation and rating period: Nociceptive stimulation began 2 seconds after the onset of the eVAS display and lasted for 6 seconds. The set of active electrodes depended on the selected trials, 5) Post stimulation period: Following the offset of stimulation, the eVAS remained visible for an additional 4 seconds and after a total of 12 seconds, the pain intensity eVAS disappeared from the screen, 6) Pain extraction rating: An eVAS assessing the ability to extract the pain sensation was displayed. Participants confirmed their rating using the trackball device, 7) Attentional focus rating: An eVAS assessing the ability to focus attention on the pain sensation was displayed. Participants confirmed their rating using the trackball device, 8) Collecting report on the location of the most intense pain intensity: Participants were asked whether stimulation from any of the electrodes was more intense than others. Participants reported the number of electrode(s). After completion of all ratings, a message reading “A short break now. Please wait patiently.” was displayed.

### 2.7. Statistical analyses

All collected raw data from each subject was first checked for completeness and quality and later pre-processed in Python3 with ‘numpy’ and ‘pandas’ libraries. Due to real-time data acquisition during stimulation periods, each rating curve was down sampled to single summary statistic: Area Under the Curve (AUC) expressing global pain level in each trial. For the purpose of data visualization raw pain curves were interpolated to a common time base and averaged per trial type within each subject and then encompassed group-level averaging **(Figure 2)**. All secondary variables including attentional, extraction and pain maxima ratings were merged with AUC values prior to performing inferential tests. Analyses were performed as outlined in the preregistered protocol, with three deviations from the protocol, for which the rationale and additional clarifications are provided in the **Appendix S1.** All statistical analyses were conducted in R Studio (R version 4.4.2; R Core Team, 2024). Linear mixed-effects models were implemented using the “lme4” package, with *p*-values obtained via “lmerTest”. Estimated marginal means and planned contrasts were calculated using the “emmeans” package with alpha level set to 0.05.

**Figure 2.**
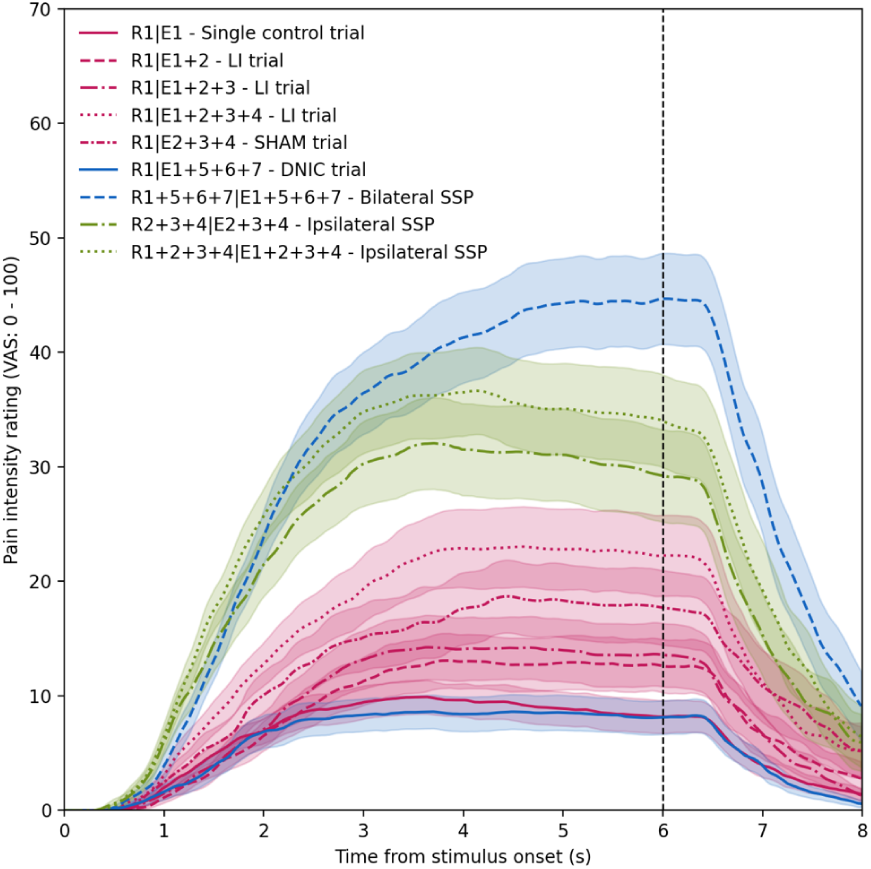
Group-level pain intensity ratings in all trials (N = 30). “R” refers to electrode from which pain was rated by the subject. “E” refers to electrode(s) which were actually active and thus stimulating in the given trial. Vertical bar points to stimulus offset. Pink coloured curves refer to lateral inhibition (LI) trials and a SHAM trial. Blue curves refer to DNIC and bilateral summation trial encompassing stimulation on the opposite arm. Green curves refer to ipsilateral spatial summation (SSP) trials. Curves represent mean pain with shaded areas representing standard errors.

#### 2.7.1. Confirmatory contrasts

To address main hypotheses outlined in the preregistration, pain AUC scores were set as dependent variable (DV) and used in the linear mixed model (LMM): “pain AUC” ∼ “trial” + (1 | “subject”) where “trial” was set as a fixed factor and “subject” was set as random intercept. More complex models were also explored including additional fixed factor “block” (3, 4) and covariate “sequence” (individual position of each trial in the sequence) and also their interactions. However, because interactions were not significant and results derived from these models were largely the same (**Appendix S9**), we retained the primary analysis. Significant main effect of “trial” was followed by *t* contrasts between selected trials and type I error was controlled by False Discovery Rate (FDR) correction applied to contrasts addressing the same hypothesis. As robustness check Šidák correction was also applied and results compared against FDR **(Appendix S8)**. As outlined in preregistration, pain from R1|E1 trial was compared to pain from R1|E1+E2, R1|E1+E2+E3 and R1|E1+E2+3+4 to address main hypothesis **(H1)** on LI. Pain from R1|E1 was contrasted with pain from SHAM trial (R1|E2+E3+E4) to address hypothesis **(H2)**. To test hypothesis **H3** that number of electrodes activated influence LI, pain from all LI trials were contrasted with each other. Pain from R1|E1 was compared to R1+R2+R3| E1+E2+E3 and R1+R2+R3+R4| E1+E2+E3+E4 to demonstrate SSP **(H4)**. To test hypothesis that stimulation over one limb with four electrodes will be more painful compared to stimulation of two limbs with four electrodes **(H5)** pain R1+R2+R3+R4| E1+E2+E3+E4 was contrasted with R1+R5+R6+R7| E1+E5+E6+E7.

#### 2.7.2. Exploratory contrasts

Because nonspecific factors such as attention, expectations, and radiation of pain may contribute to the inhibitory mechanisms, we sought to extract the pain response during the R1|E1+E2+E3+E4 trial while accounting for these nonspecific influences. To this end, a subject-level effect - the difference in AUC between the R1|E1+E2+E3+E4 trial and the sham trial (R1|E2+E3+E4) - was calculated first and then compared with the baseline pain produced by the R1|E1 trial. Further exploratory contrasts involved comparison between R1|E1 and R1|E1+E5+E6+E7 to test for the presence of significant DNIC effect, and also between R1|E1+E5+E6+E7 and R1+R5+R6+R7|E1+E5+E6+E7 and between R1|E1+E2+E3+E4 and R1+R2+R3+R4|E1+E2+E3+E4 to test for attentional modulation of pain. To test if effect from main LI trial (R1|E1+E2+E3+E4) is distinct from DNIC it was contrasted with (R1|E1+E5+E6+E7). All these five contrasts were FDR controlled under one family of tests.

#### 2.7.3. Additional exploratory analyses

To characterize attribution of pain maxima to each electrode, we estimated the probability of selecting given electrode under each trial. Localisation responses were treated as Bernoulli trials (selected vs not selected), restricted to identifiable responses (i.e., excluding “no” responses). For each trial, responses were pooled across blocks 3 and 4. Probability estimates were obtained using a Beta-Binomial model with a uniform Beta_(1,1)_ prior, yielding posterior distributions over the probability of selecting each electrode separately. From these distributions, posterior means and 95% credible intervals (CrI) were computed. To investigate the role of pain maxima attribution in the context of LI, comparisons were narrowed down to critical trials that involved ratings only from E1: (R1|E1, R1|E1+2+3+4, SHAM: R1|E2+3+4 and DNIC: R1|E1+5+6+7). Posterior distributions of probability differences (ΔP) were generated by Monte Carlo sampling (60,000 draws per condition), allowing direct estimation of the probability that ΔP exceeded zero. Because this analysis revealed highly significant drop in probability of selecting E1 as a source of pain maxima, the follow-up contrasts were made by analysing pain AUC values in trials of interest (R1|E1+2+3+4, and SHAM: R1|E2+E3+E4) and dividing subjects in those who attributed pain maxima to middle (rated) electrode and these who experienced maxima in other electrode(s) not including E1.

To assess whether cognitive and perceptual factors moderated the observed facilitatory effect, the subject-level LI effect was calculated as (Δ) difference between R1|E1+2+3+4 and R1|E1 with positive values indicating facilitation and negative values indicating inhibitory responses. Such a new subject-level metric (ΔLI) was tested for the relationship with the following predictors: i) non-specific pain AUC observed in the SHAM trial, ii) the attentional drift (difference in VAS attention ratings), iii) pain extraction drift (difference in VAS extraction ratings), and iv) baseline sensitivity as reflected in pain AUC scores in R1|E1 trial. Importantly, VAS-derived attention and extraction ratings indexed target-specific processes, with <u>lower</u> Δ values suggesting attention (or pain extraction) drop in R1|E1+2+3+4 compared to R1|E1. In other words, Δ values close to “0” would suggest constant attention/extraction extent across these two trials. All these regressions were tested for moderator effect of pain maxima attribution (E1 vs non-E1).

## 3. RESULTS

### 3.1. Basic characterization, quality check, and main model

Data from two participants were excluded due to no pain reports at the intensity of stimuli used in the study, and the data from the first participant was excluded from the analysis, as the first administration of the procedure was conducted for experimenter training. The remaining participants successfully completed all study procedures. Thus, data from a total number of 30 participants was used in statistical analyses, in line with the protocol. Descriptive statistics are presented in **Appendix S2**.

Data from the first block of stimuli showed that sensitivity of skin sites the electrodes were attached to were largely comparable with lack of significant differences in pain AUC in 5 out of 6 comparisons (see **Appendix S6)**. Furthermore, pain-site localization was highly accurate: 87% of participants correctly identified E1, 90% E3 and E4, and 93% correctly identified electrode E2. The main LMM analysis performed on data from blocks 3 and 4 revealed significant effect of “trial” (*F*_(8, 232)_ = 35.23, *p* < 0.001, *η_p_^2^* = 0.55), which is decomposed into contrasts of interest in the following sections and shown on **Figure 3**.

**Figure 3.**
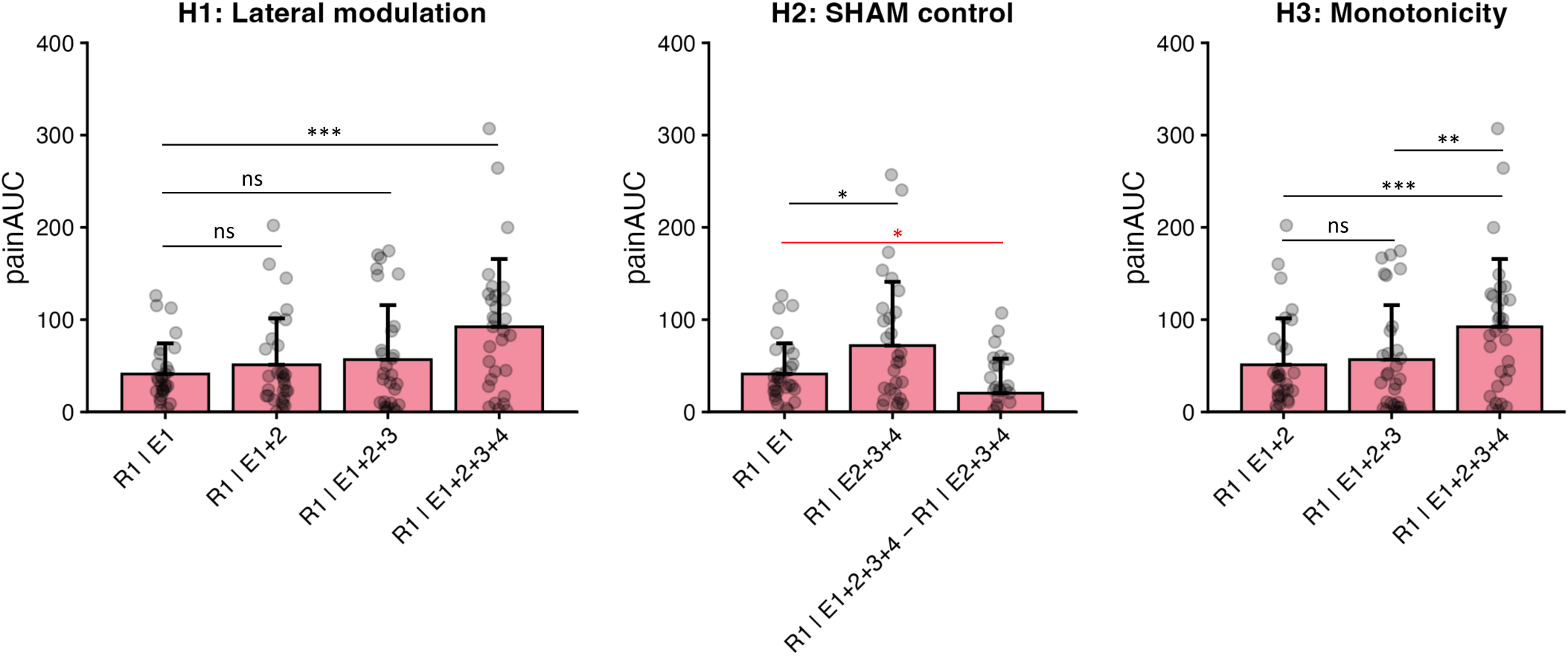
Primary analyses. Hypothesis H1: Direct contrasts between lateral inhibition (LI) trials and single electrode trials (R1|E1) did not show inhibition. Instead, pain from a target electrode (E1) increased when three adjacent electrodes (E2, E3, and E4) were co-activated. H2: Pain in a SHAM trial was substantial despite no stimulation applied to E1 from which pain was rated. Follow-up comparison revealed that significant portion of pain from E1 electrode during R1|E1+2+3+4 is caused by the pain radiation from adjacent electrodes: Exploratory contrast between R1|E1 and subject-level cleaned inhibitory response (R1|E1+2+3+4 minus SHAM) was significant. H3: A monotonic increase in spatial input led to a significant increase in pain only after a critical threshold of three additional electrodes was activated. Plots depict mean values ± standard deviations. Red line shows exploratory contrast. \**p*_FDR_ < 0.05, \*\**p*_FDR_ < 0.01, \*\*\**p*_FDR_ < 0.001.

### 3.2. Lateral inhibition and facilitatory effects

**Hypothesis (H) 1:** Primary contrasts were conducted to test the hypothesis that pain elicited at E1 is reduced when accompanied by concurrent co-stimulation of adjacent electrodes (**Figure 3**). Compared with R1|E1, stimulation with one (R1|E1+E2) or two (R1|E1+E2+E3) additional electrodes did not lead to statistically significant decreases in perceived pain at the target site (R1|E1 vs. R1|E1+E2: *M* = 41.16 vs. 51.15, *t*_(232)_ = −0.82, *p*_FDR_ = 0.42, *d*_z_ = −0.21; R1|E1 vs. R1|E1+E2+E3: *M* = 41.16 vs. 56.76, *t*_(232)_ = −1.28, *p*_FDR_ = 0.31, *d*_z_ = −0.24). In contrast to our hypothesis, stimulation with three neighbouring electrodes (R1|E1+E2+E3+E4) produced a robust and statistically significant increase rather than decrease in perceived pain at the target electrode relative to R1|E1 trial (*M* = 41.16 vs. 92.26), reflected in a large within-subject effect (*t*_(232)_ = −4.18, *p*_FDR_ < 0.001, *d*_z_ = −0.72). Thus, H1 was not supported since data revealed facilitatory instead of inhibitory effects.

**H2:** To address the hypothesis that pain originating from SHAM stimulation is negligible (i.e., lower than in the R1|E1 trial), and to control for this potential confound, we contrasted the pain AUC in the R1|E1 condition (target electrode stimulated alone) with the SHAM trial (R1|E2+E3+E4) (**Figure 3**). In the SHAM trial, only the surrounding electrodes were activated, while participants rated pain at the unstimulated target electrode (E1). Contradictory to expected outcome, pain AUC was significantly higher in the sham trial (R1|E2+E3+E4) compared to R1|E1 trial (*M* = 71.90 vs. 41.16), yielding a moderate within-subject effect (*t*_(232)_ = 2.51, *p*_FDR_ < 0.05, *d*_z_ = −0.45). This result may suggest that potential pain reductions in the main LI trial (R1|E1+2+3+4) are overshadowed by pain radiating from adjacent electrodes. Exploratory contrasts provide further insight into the mechanisms contributing to these effects. To isolate pain attributable specifically to active target stimulation beyond nonspecific spatial contributions, a subject-level effect, i.e. difference in AUC between R1|E1+E2+E3+E4 and SHAM: R1|E2+E3+E4 was calculated first. Then it was compared with baseline pain produced by R1|E1 trial. Residual pain following SHAM subtraction was significantly lower than pain during R1|E1 (*M* = -20.80, *t*_(29)_ = −2.33, *p* < 0.05, *d*_z_ = −0.43), indicating that a substantial portion of the pain observed in the main LI trial (R1|E1+E2+E3+E4) reflects nonspecific spatial or contextual effects rather than direct nociceptive input from the target electrode alone.

**H3:** To verify hypothesis that the spatial load from adjacent electrodes had monotonic effect on reported pain from a target, pain AUC was compared across LI trials **(Figure 3)**. While pain from R1|E1+E2 trial was not different from R1|E1+E2+E3 (*t*_(232)_ = −0.46, *p*_FDR_ = 0.65, *d*_z_ = −0.12,), the pain increased in R1|E1+E2+E3+E4 compared to R1|E1+E2 (*t*_(232)_ = −3.36, *p*_FDR_ < 0.01, *d*_z_ = −0.70,) and R1|E1+E2+E3 trial (*t*_(232)_ = −2.90, *p*_FDR_ < 0.01, *d*_z_ = −0.76). Together, these findings indicate that pain was greatest when all adjacent electrodes were co-activated.

### 3.3. Spatial summation effects

**H4:** To examine the SSP effects, single-electrode stimulation (trial R1|E1) was compared with trials involving stimulation of multiple electrodes and overall pain ratings (see **Figure 1** for details) under the hypothesis that summation is present if more intense pain is observed in trials with greater number of electrodes activated **(Figure 4)**. In fact, pain AUC increased markedly as the spatial extent of stimulation increased: Relative to R1|E1 (*M* = 41.16), pain was substantially higher in R1+R2+R3|E1+E2+E3 (*M* = 134.67), yielding a large within-subject effect (*t*_(232)_ = −7.64, *p*_FDR_ < 0.001, *d*_z_ = −1.11). This effect was even more pronounced for R1+R2+R3+R4|E1+E2+E3+E4 (*M* = 155.39), resulting in a larger effect size (*t*_(232)_ = −9.33, *p*_FDR_ < 0.001, *d*_z_ = −1.46). These findings demonstrate robust SSP, with pain intensity scaling strongly with the number of simultaneously activated nociceptive sites.

**Figure 4.**
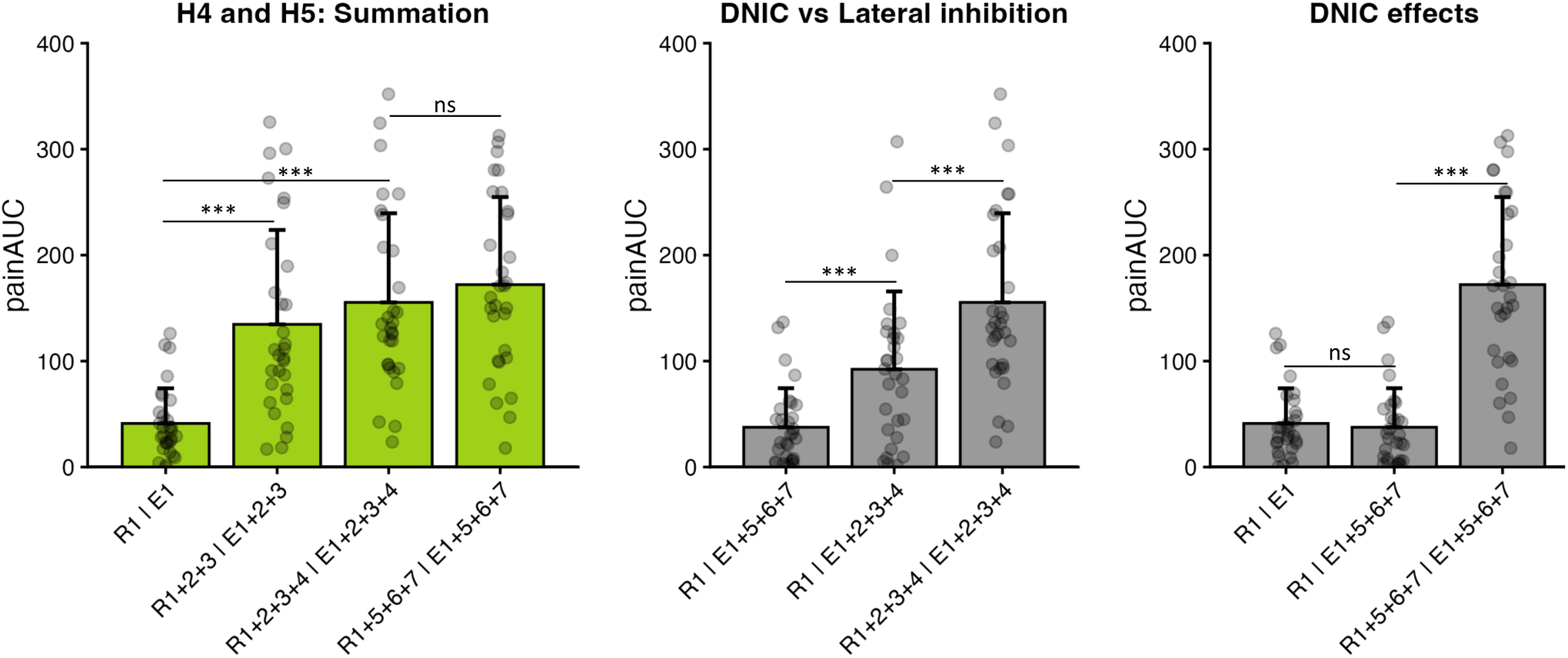
Secondary analyses. The left panel illustrates hypotheses H4 and H5. Ipsilateral spatial summation of pain was significant: increasing the number of active electrodes resulted in higher pain intensity. In contrast, no significant difference was observed between R1+2+3+4|E1+2+3+4 and R1+5+6+7|E1+5+6+7. Exploratory analyses are presented in the middle and right panels (grey bars). Pain at E1 was significantly reduced during the diffuse noxious inhibitory control (DNIC) trial (R1|E1+5+6+7) compared with the main lateral inhibition (LI) trial (R1|E1+2+3+4). Spatial summation was abolished under the directed attention rating procedure, as noted by significant difference between R1|E1+2+3+4 and R1+2+3+4|E1+2+3+4, and between R1|E1+5+6+7 and R1+5+6+7|E1+5+6+7. The DNIC trial did not significantly reduce pain relative to baseline (R1|E1). Plots depict mean values ± standard deviations. \**p*_FDR_ < 0.05, \*\**p*_FDR_ < 0.01, \*\*\**p*_FDR_ < 0.001.

**H5:** To address the hypothesis that SSP is present ipsilaterally but not bilaterally, pain AUC from R1+R2+R3+R4|E1+E2+E3+E4 was compared with R1+R5+R6+R7|E1+E5+E6+E7 **(Figure 4)**. Pain AUC values were slightly lower for ipsilateral than bilateral summation (*M* = 155.39 vs 172.16), although the difference did not reach statistical significance (*t*_(232)_ = −1.37, *p*_FDR_ = 0.17, *d*_z_ = −0.22). This result suggests not only unilateral but also bilateral SSP and trend towards higher pain in bilateral stimulation.

### 3.4. Exploratory contrasts and DNIC effects

The classic approach of demonstrating the DNIC effect did not show a significant result **(Figure 4**): Pain AUC in the R1|E1 trial was not different from pain in R1|E1+E5+E6+E7 trial (*t*_(232)_ = −0.29, *p* = 0.77, *d*_z_ = −0.12) suggesting no evidence of a DNIC effect under the present experimental conditions. To evaluate whether relocating the spatial stimulation from the ipsilateral to the contralateral side engaged DNIC-related inhibitory mechanisms, we first compared pain from R1|E1+E5+E6+E7 trial (DNIC) with main LI trial R1|E1+E2+E3+E4 **(Figure 4)**. However, pain significantly increased during the main LI trial (*M* = 92.26) than in the DNIC (*M* = 37.55), yielding a large effect (*t*_(232)_ = −4.47, *p*_FDR_ < 0.001, *d*_z_ = −0.84). Together, these results may suggest that relocating spatial load reduced facilitatory influences seen in ipsilateral stimulation, but within current protocol did not produce significant DNIC modulation.

Interestingly, pain in R1|E1+E5+E6+E7 (*M* = 37.55) was significantly lower compared to bilateral summation trial R1+R5+R6+R7|E1+E5+E6+E7 (*M* = 172.16), in which the same configuration of electrodes was activated, resulting in large effect size (*t*_(232)_ = −11.00, *p* < 0.001, *d*_z_ = −1.76). Consistent with this, overall pain in main LI trial R1|E1+E2+E3+E4 was significantly lower than during R1+R2+R3+R4|E1+E2+E3+E4 trial (*M* = 155.39 vs. 92.26, *t*_(232)_ = −5.16, *p* < 0.001, *d*_z_ = −1.02). These results demonstrate the role of top-down mechanisms which can, accordingly, reduce or enhance SSP.

### 3.5. Exploration of drifts in localisation of pain maxima

In general, participants’ selection of electrodes producing pain maxima (pain of highest intensity) distributed differently according to trial type **(Figure 5A and B)**. Posterior means and 95% credible intervals (CrI) showed near-ceiling localisation in R1|E1 (mean = 0.96, 95% CrI [0.90, 1.00]) but substantially reduced localisation in R1|E1+2+3+4 (0.40 [0.27, 0.54]) and sham trial R1|E2+3+4 (0.31 [0.19, 0.45]). Posterior contrasts confirmed that localisation probability of selecting the middle electrode E1 as the source of pain maxima was markedly lower in R1|E1+2+3+4 relative to R1|E1 (posterior probability = 0.99, mean diff, = -0.56, 95% CrI [-0.70, -0.41]). Localization of E1 was lowest in the DNIC trial R1|E1+5+6+7 relative to R1|E1 (posterior probability = 0.99, difference = −0.95, 95% CrI [−0.99, −0.87]), whereas R1|E2+3+4 and R1|E1+2+3+4 did not differ (posterior probability = 0.18, CrI including zero) **(Figure 5C)**.

**Figure 5.**
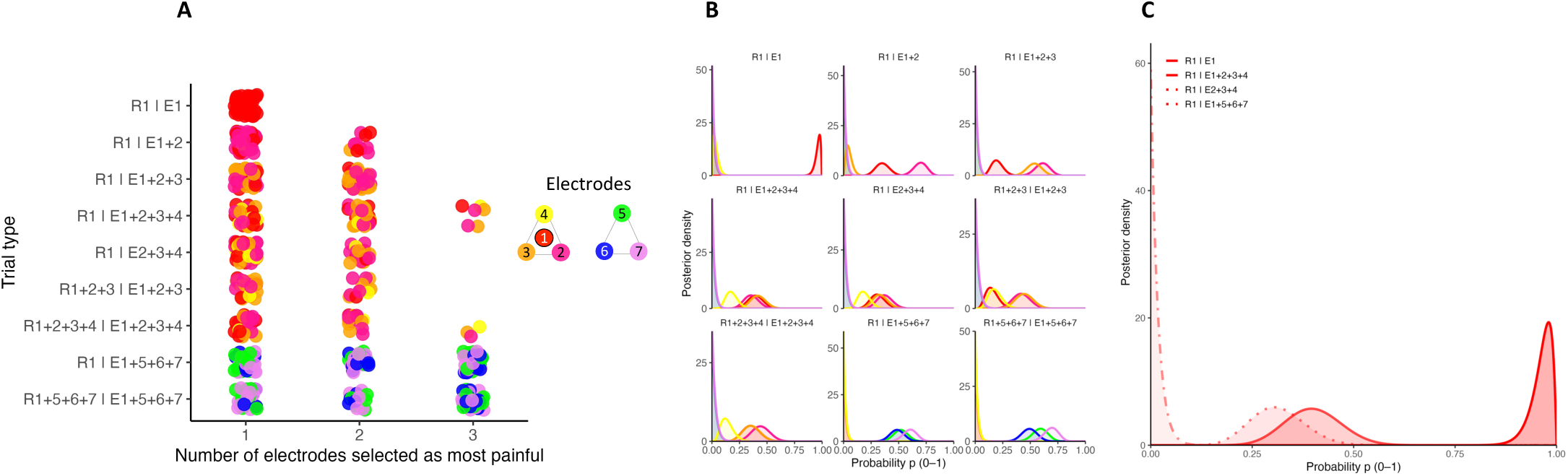
Pain maxima localization(s) across trials. (A) Pain maxima ratings given by all subjects and trials from the main phase of the experiment (every dot represents a single rating). Red cluster of responses is noted for R1|E1 trial in which only middle electrode (E1) was activated. In general, adding more electrodes to the system showed that participants’ responses were more diverse, often involving more than one electrode as a source of pain maxima (lower axis shows amount of electrodes picked as a source of pain maxima: from one to three). (B) Posterior probability density functions for selecting each electrode in all trials. Densities were derived from a Beta–binomial model with a uniform prior, using pooled block-level localization responses from blocks 3 and 4 and excluding non-identifiable responses (“no”). (C) Posterior probability density functions for selecting E1 in critical trials: LI0 (R1|E1), LI3 (R1|E1+2+3+4), SHAM (R1|E2+3+4), DNIC (R1|E1+5+6+7). Separation between curves illustrate systematic shifts in pain maxima attribution across these trials.

Attributing pain maxima to the target E1 was associated with changes in pain intensity. Exploratory LMM analysis was run using trials of relevance (R1|E1+2, R1|E1+2+3, R1|E1+2+3+4, and SHAM: R1|E2+E3+E4), trial-specific attribution (selection of E1), and a random intercept for participant: “painAUC” ∼ “trial” × “E1 selection” + (1 | subject). The model revealed significant main effects of “trial” (*F*_(3, 79.73)_ = 8.23, *p* < 0.001, *η_p_^2^* = 0.24) and “E1 selection” (*F*_(1, 93.09)_ = 16.47, *p* < 0.001, *η_p_^2^* = 0.15), indicating that pain AUC increased both with spatial extent of stimulation and with attribution of pain to the E1. In contrast, the “trial” × “E1 selection” interaction was not significant (*F*_(3, 82.48)_ = 0.37, *p* = 0.77, *η_p_^2^* = 0.01), suggesting that the effect of attribution on pain magnitude was comparable across trials. Overall, participants who attributed maxima to E1 had significantly higher pain AUC compared to participants who felt maxima outside of the target **(Figure 6)**.

**Figure 6.**
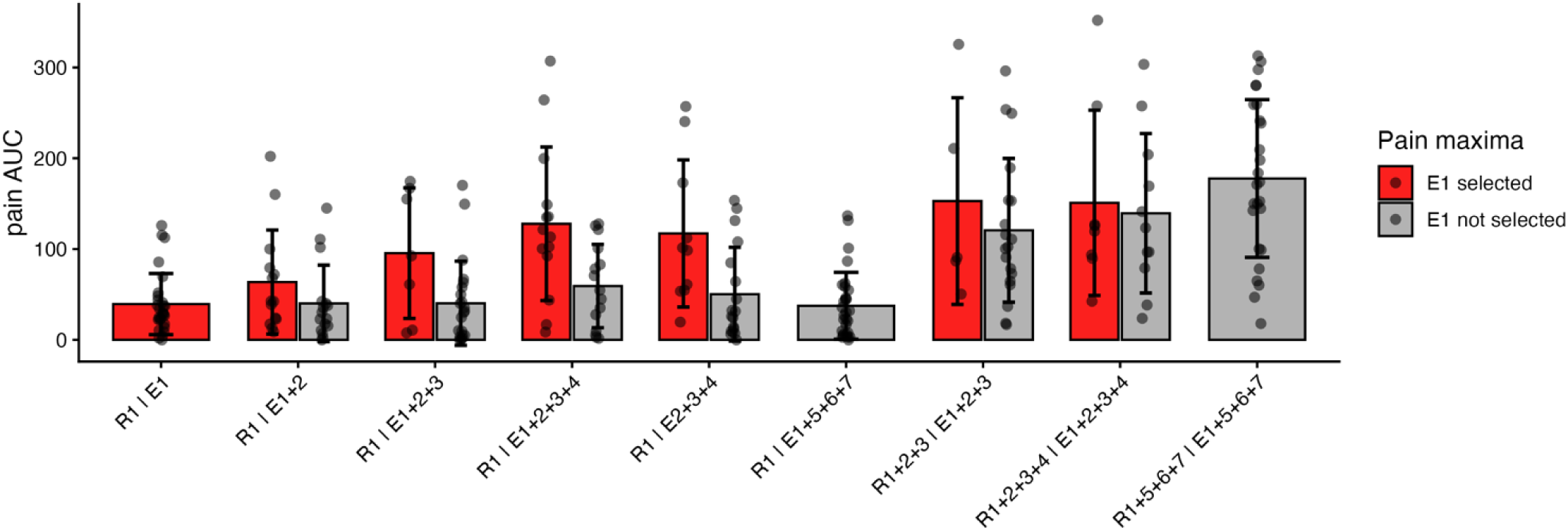
Pain AUC obtained from a target E1 electrode depends on pain maxima localization. Participants in R1|E1 trial selected E1 as a source of pain in 28/30 cases (two subjects did not point to particular electrode). In lateral inhibition LI1 (R1|E1+2), LI2 (R1|E1+2+3), LI3 (R1|E1+2+3+4) trials and the SHAM trial (R1|E2+3+4) those subjects who reported pain maximum in the E1 also experienced the highest pain.

### 3.6. Moderating role of pain maxima locus

To explore whether cognitive and perceptual factors moderated the observed effects in main LI trial, the subject-level LI effect was calculated and its relationship investigated with the following predictors: i) non-specific pain AUC observed in the sham trial, ii) the attentional drift (difference in attention ratings), iii) pain extraction drift (difference in extraction ratings), and iv) baseline sensitivity as reflected in pain AUC scores in R1|E1 trials. Importantly, VAS-derived attention and extraction ratings indexed target-specific processes, with higher values indicating greater attentional focus on the target E1, or perceptual extraction of pain from E1. Magnitude of pain AUC in SHAM vs. LI effect showed a trend-level interaction with “E1 selection” subgroup (*p* = 0.07, **Figure 7A**). Simple-slope analyses revealed pain in SHAM trial robustly predicted subsequent pain amplification (LI effect) in participants who attributed pain maxima to E1 during R1|E1+2+3+4 trial (*β* = 0.83, *p* < 0.001), whereas no such relationship was observed in participants who did not (*β* = 0.33, *p* = 0.15). Critically, changes in target-directed attention (ΔVAS-Attention) significantly interacted with “E1 selection” grouping (*p* < 0.05, **Figure 7B**). In the E1-subgroup, increases in attention toward the target electrode were associated with greater pain amplification (*β* = 2.16, *p* < 0.05). In contrast, no relationship was observed in the subgroup who did not attribute maxima to E1 (*β* = −0.20, *p* = 0.77). Negative values of ΔVAS-Attention, reflecting loss of attentional focus on the target electrode during E1|R1+2+3+4, were associated with attenuated pain amplification in the E1-subgroup. In contrast, changes in the perceived ability to extract pain from the target electrode (ΔVAS-extraction) did not interact with perceptual grouping (*p* = 0.23), and simple slopes were non-significant in both subgroups (**Figure 7C**). This dissociation suggests that attentional allocation to the target electrode, rather than introspective clarity of pain origin, modulates pain facilitation. No moderation was found between baseline sensitivity and LI effect nor their interaction with pain maxima strata (**Figure 7D**).

**Figure 7.**
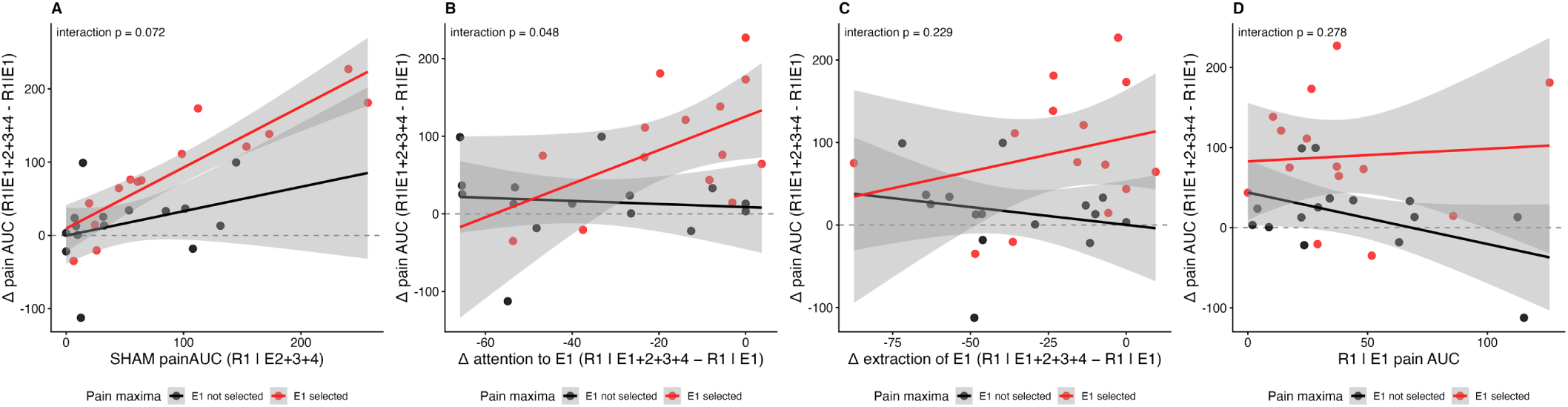
Moderation analyses of lateral inhibition/facilitation effects. The moderating role of selecting electrode E1 as the source of the pain maximum was explored across a series of theoretically plausible relationships. (A) No significant interaction was observed between LI effects (difference in pain AUC for R1|E1+2+3+4 vs. R1|E1) and pain reported during the SHAM trial (R1|E2+3+4), in which substantial pain was perceived under a non-stimulated electrode. (B) When participants identified E1 as the location of the pain maximum, the LI effect (with positive values indicating facilitation) was positively associated with the attentional drop index (the difference in VAS attention ratings between R1|E1+2+3+4 and R1|E1). (C) No relationship was observed between LI effects and extraction ratings (ability to extract target pain from pain induced by adjacent electrodes). (D) LI effects were not related to a baseline sensitivity as assessed by pain AUC in R1|E1 trial.

## 4. DISCUSSION

Contrary to our original predictions, the designed LI paradigm used in the current study revealed facilitatory rather than inhibitory profiles. Control analyses further revealed that substantial pain was reported even during SHAM trial, uncovering pronounced radiation [4,57] and SSP phenomena [13,15,25,53]. Pain ratings were highest when all adjacent electrodes were co-activated, with marked facilitation emerging only beyond a critical spatial extent of nociceptive input. Changes in attention critically contributed to the observed facilitation and this effect was moderated by the perceived spatial localization of the pain maximum. Facilitation emerged when spatial nociceptive input expanded and participants maintained attention on the target (rated) electrode, with the pain maximum accurately attributed to that location. In contrast, attention-driven facilitation was absent when the pain maximum was perceived outside the target electrode.

These findings are consistent with the possibility that facilitatory effects of spatially expanded nociceptive input depend on attentional focus and the spatial attribution of pain maxima, although the present design does not allow these mechanisms to be isolated directly. Lastly, the paradigm revealed for the first time existence of SSP induced through nociceptive input from homologues contralateral sites of the body. Because the present study relied exclusively on psychophysical measures, the underlying neural mechanisms cannot be directly inferred. Thus, the proposed interpretations should be considered as plausible mechanistic explanations.

### 4.1. Lateral facilitation in nociceptive processing

Capturing LI in vivo in humans entails methodological constraints, as direct recording of cellular-level neuronal changes while experimentally manipulating the microscale environment is not feasible in awake human participants. Nevertheless, careful translational inference from neuronal [23,44] and animal models [24] to human research [20,58] allows the formulation of plausible assumptions that may, at least in part, account for the present findings. Interestingly, using the current LI paradigm pain ratings were not reduced. Instead, they remained stable until all three adjacent electrodes were co-activated, at which point pain increased significantly. The observed facilitation suggests that inhibitory interactions predicted by the LI model may be -in presented conditions-overcome once nociceptive input exceeds a critical spatial threshold. In the present study, two facilitatory processes were psychophysically evident: SSP [5,40,53] and radiation of pain [4,55]. Increasing the spatial load of nociceptive stimulation may recruit adjacent functional receptive fields (RFs) whose activation thresholds are reached only when multiple spatial foci are engaged simultaneously. Such widespread recruitment may also give rise to radiation phenomena, whereby pain is perceived beyond the area of direct nociceptive input [4]. Consistent with this interpretation, participants reported substantial pain in the SHAM condition despite the absence of direct stimulation at the rated site, suggesting that radiation and potentially expectancy effects contributed to the reported sensations. Together, these facilitatory processes may have overridden inhibitory interactions predicted by the LI framework. Consequently, our attempt to isolate “pure” LI previously demonstrated in paradigms with noxious [20,58] and innocuous heat [31], may have been masked by stronger facilitatory mechanisms operating at the population coding level. Such facilitatory profiles may partly reflect differences in stimulus parameters, including the sustained nociceptive input used in the present study compared with the shorter stimuli employed in previous LI paradigms [3,20,25]. Alternatively, the intensity of nociceptive input in the present study may have been insufficient to engage robust inhibitory mechanisms [3].

### 4.2. Lateral inhibition in nociceptive processing

Interestingly, not all participants exhibited facilitatory profiles. Secondary measures, specifically the perceived localization of the pain maxima and the extent to which participants maintained attention on the target allowed for additional inferences. As could be predicted, participants consistently reported the maximum pain as originating from the target site in trials in which only the target electrode (E1) was active. However, the probability of localizing the pain maximum at the target substantially decreased in LI trials, in which participants were more likely to report pain maxima at a non-target site. In the main LI trial (R1|E1+2+3+4), approximately half of participants reported that the pain maximum was located outside the target site. Importantly, these participants experienced significantly lower pain compared with those who reported the pain maximum at the target site. Exploratory moderation analyses revealed that attentional focus played a significant role in shaping these facilitatory profiles. Specifically, facilitation was observed only in participants who reported the pain maximum at the target and maintained stable attention on it. When attentional focus declined, facilitation was abolished, particularly in participants who otherwise identified the target electrode as the source of maximal pain. Abolished facilitation was maximized in DNIC trial, in which three electrodes were active yet repositioned to the contralateral side.

Any inhibitory component may have been further modulated by attentional drift towards more salient nociceptive foci. Such shifts of attention could alter the effective population of activated neurons, potentially increasing neuronal recruitment while narrowing the RFs contributing to the percept, thereby reducing spatial summation. Supporting this interpretation, previous work showed that line-like stimulation can evoke less pain than two-point stimulation despite covering a larger area, although responses were highly variable [58], suggesting that inhibitory interactions are sensitive to contextual factors. Behavioral studies further demonstrate that directing attention to a specific part of a nociceptive foci reduces reported pain [2,56], whereas divided attention –requiring dynamic switching between nearby nociceptive foci– increases the magnitude of inhibition [2,12,56]. Neuroimaging studies similarly indicate that reduced attention to nociceptive input is associated with lower pain ratings, increased engagement of descending modulatory regions [7,51,66,68], and reduced activity at the spinal cord [64]. Evidence from DNIC studies also indicates that attention can attenuate inhibitory interactions: participants report greater pain when attending to the rated stimulus than when focusing on a competing nociceptive input [14,32,61].

### 4.3. Lateral modulation vs. summation and DNIC

Although increased overall pain with greater spatial extent of nociceptive stimulation was anticipated –and is consistent with previous work from our group [3,5,45] and others [40,53]– the presence of spatial summation when stimulation was applied bilaterally to opposite limbs far from the midline is novel. A reduction in pain was expected under bilateral stimulation because of the potential contribution from DNIC mechanisms. Unlike classical DNIC paradigms [14,19,22,60], which use a sustained conditioning stimulus that precedes the test stimulus, the present study used brief conditioning and test stimuli that were applied simultaneously. Such temporal dependency may have limited the recruitment of descending inhibitory circuits and contributed to the absence of DNIC. Observation of bilateral SSP extends current models of summation, which have been derived primarily from within-[17,46,57] and across-dermatomal [17,33,46,57] configuration of stimulating probes. Notably, an earlier study by Defrin and colleagues [14] using two heat probes placed on opposite forearms, reported pain ratings comparable to single-site stimulation – a finding that provided support for our original prediction. Whether observed summation was a result of using different modality and underlying activation of divergent peripheral apparatus is not clear and must be further investigated. Nevertheless, the observed bilateral summation –and its attentional tuning [2,56], whereby stimulation geometry was held constant while ratings were anchored to a target unilateral site– points to a predominantly central mechanism of SSP. Single-unit electrophysiological data indicates that much SSP could take place in the spinal cord. Nociceptive neurons within the deep dorsal horn and parts of the ventral horn have large and even bilateral RFs that could subserve widespread spatial summation [21]. Propriospinal interconnections [11,49,50,63] could easily support such spatial integration across dermatomes and the body midline.

### 4.4. Spatial-attentional regulation of pain

Spatial integration of nociceptive input may depend not only on the physical extent of stimulation but also on attentional allocation and on the perceived locus of maximal pain. Under such conditions, LI may be masked by facilitatory mechanisms such as spatial summation. This framework may explain why attentional manipulations during nociceptive stimulation produce both analgesic and hyperalgesic effects. Small shifts of attention across spatially distributed inputs may determine whether facilitatory or inhibitory interactions dominate. Attentional control of spatial integration may therefore represent a key mechanism shaping pain perception. Taken together, the present findings may primarily reflect an interaction between spatial summation and cognitive factors, particularly attentional allocation, whereas the contribution of LI mechanisms cannot be determined directly from the present psychophysical data.

### 4.5. Limitations

First, the stimulation paradigm was not designed to selectively activate small-diameter nociceptive C- and Aδ-fibers. Instead, the electrical stimulation with rectangular pulses likely co-activated large-diameter Aβ fibers [27,65], meaning that neuromodulatory mechanisms associated with mixed afferent input, including gate-control mechanisms [28,43], may have contributed to the observed pain responses. Although this limits conclusions regarding specific afferent contributions, identical stimulation parameters across all conditions reduce concerns about effects on between-condition comparisons. Second, although designed to maximize LI, the present paradigm did not fully isolate LI from other concurrent mechanisms, including summation, radiation, and attention. Therefore, conclusions regarding the balance between inhibitory and facilitatory nociceptive mechanisms should be interpreted with appropriate caution. Future studies using selective nociceptive stimulation and paradigms disentangling these mechanisms in causal fashion in healthy and clinical populations are needed to better understand spatial aspects of pain modulation.

## ACKNOWLEDGEMENTS

The authors have no conflicts of interest to declare. Language editing during writing process was supported by the AI tool (ChatGPT) and components of the design figure were generated by AI. The study was funded by the National Science Centre (NCN), Poland (2020/37/B/HS6/04196). WMA is supported by the National Science Centre in Poland under grant no. 2025/57/B/HS6/01722 and JS is supported by grant no. 2020/37/N/HS6/04210. Data and code supporting the findings are available upon reasonable request and will be publicly released in a repository upon publication.

