## Appendix S1-S9 for "Interaction of attention and localization of pain maxima shift the balance between lateral inhibition and spatial facilitation in nociceptive processing"

**Appendix (supplementary materials)**

### S1. Transparency statements and deviations from the preregistration

At the time of preregistration, data from 20 participants had already been collected. However, no data processing or analysis had yet been performed. Analyses were performed as outlined in the preregistered protocol, with the following deviations and clarifications. First, H2 was tested using the contrast specified in the protocol (R1|E1 vs. SHAM). This approach was retained because the wording of the preregistered hypothesis did not fully correspond to the contrast specified in the analysis plan, where SHAM was compared with R1|E1 rather than with the main LI trial. Second, two comparisons listed under H4 were omitted because they did not directly test the spatial summation hypothesis, as they involved ratings obtained under directed-attention conditions (R1|E1+E2 and R1|E1+E2+E3 vs. SSPi and SSPb). Finally, because the primary analyses revealed unexpected findings, additional exploratory analyses were conducted to examine potential factors associated with the observed pattern of facilitation. These analyses are explicitly reported as exploratory, and no confirmatory claims are based on them. In addition, for readability, target Electrode no. 4 from the preregistered protocol is referred to as Electrode 1 in the manuscript. This change was purely terminological and did not affect the experimental procedure, analyses, or interpretation of the results.

### S2. Descriptive characteristics of the study population

| Variable | Mean (SD) |
|---|---|
| Age (years) | 23.60 (4.30) |
| Fear of pain (NRS 0-10) | 2.17 (1.98) |
| FPQ-III total score | 67.10 (16.74) |
| Body mass (kg) | 75.00 (16.28) |
| Height (cm) | 175.42 (10.96) |
| BMI | 24.14 (3.49) |
| **Variable** | **N** |
| Sex | F = 14, M = 16 |
| Handedness | R = 28 (93.33%), L = 2 (6.67%) |
| Orientation of the triangle's apex | P = 17 (56.67%), D = 13 (43.33%) |

Note: F- Female, M -Male, R - right, L – left, NRS - numeric rating scale, FPQ-III – Fear of pain questionnaire, BMI- body mass index, SD – standard deviation, D – distal orientation of the triangle's apex, P – proximal orientation of the triangle's apex.

### S3. Responses to control questions following completion of the experimental procedure

| | Question | Answers (%) |
|---|---|---|
| 1 | Did you ever feel during stimulation that the location of the most intense pain shifted over time between electrodes? (yes/no) | yes = 13 (43.33%)<br>no = 17 (56.67%) |
| 2 | Did you ever feel during stimulation that the location of most intense pain was between two or three electrodes? (yes/no) | yes = 19 (63.33%)<br>no = 11 (36.67%) |
| 3 | What do you think was the goal of this study? (open) | correct = 0 (0.00%)<br>partially correct = 2 (6.67%)<br>incorrect = 28 (93.33%) |

Note: Responses to Question 3 (open-ended): correct (exact study purpose), partially correct (mentioned pain inhibition or pain summation), incorrect (did not mention pain modulation at all).

### S4. Exploratory correlational analyses

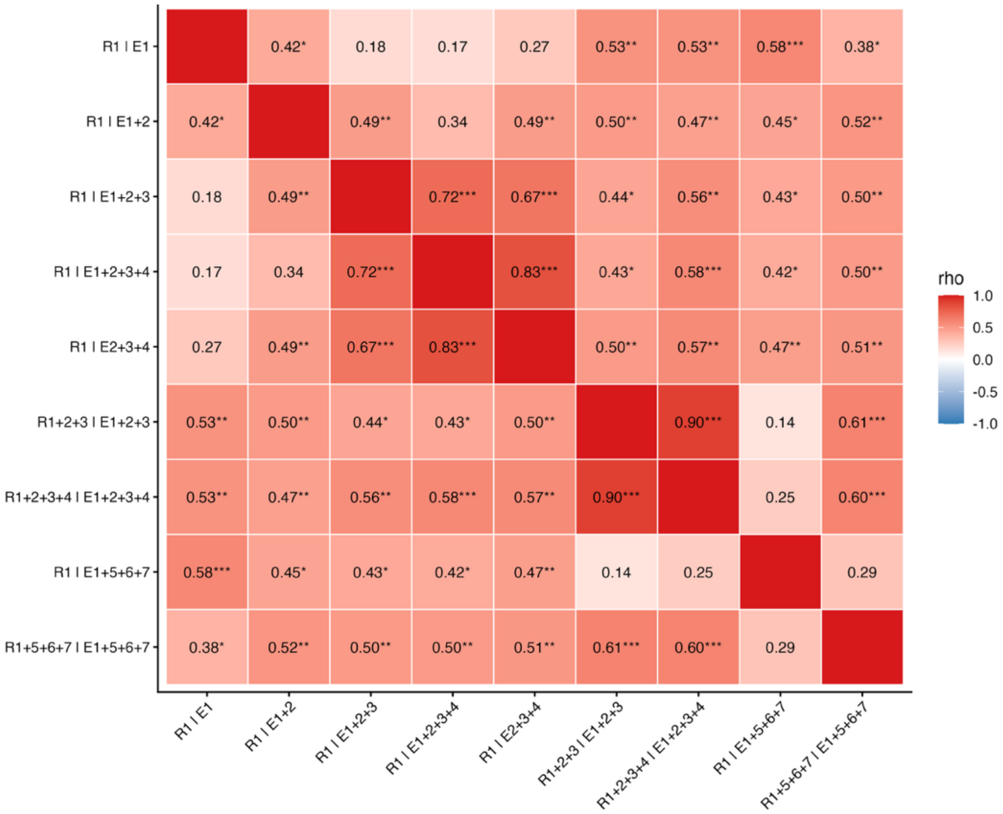


Note: Heatmap with Spearman coefficients ($r_s$) between pain AUC values from all trials.*p < 0.05, **p < 0.01, ***p < 0.001.

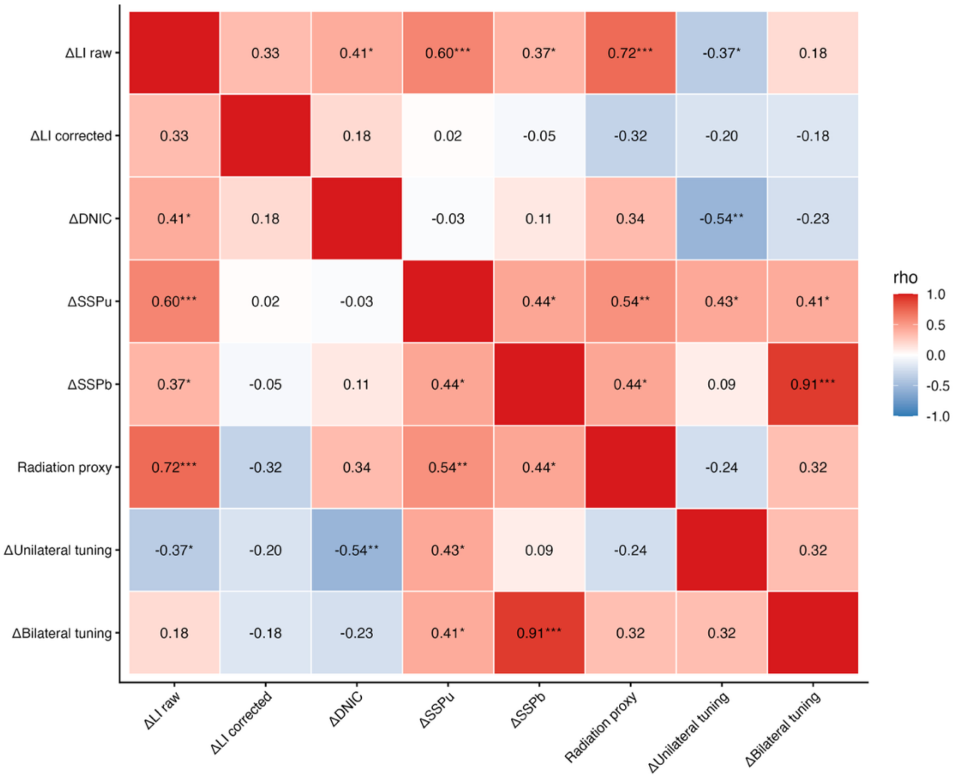


**Note:** Heatmap with Spearman coefficients ($r_s$) between individual within-subject effects, here presented with delta (Δ) sign: ΔLI raw → difference between R1|E1 and R1|E1+E2+E3+E4, ΔLI corrected → difference between R1|E1 and (R1|E1+E2+E3+E4 minus R1|E2+E3+E4), ΔDNIC → difference between R1|E1 and R1|E1+E5+E6+E7, $\Delta SSP_u$ → difference between R1|E1 and R1+R2+R3+R4|E1+E2+E3+E4, $\Delta SSP_b$ → difference between R1|E1 and R1+R5+R6+R7|E1+E5+E6+E7, Radiation proxy → pain AUC in "SHAM" trial R1|E2+E3+E4, Unilateral tuning → difference between R1+R5+R6+R7|E1+E5+E6+E7 and R1|E1+E2+E3+E4. Bilateral tuning → difference between R1+R5+R6+R7|E1+E5+E6+E7 and R1|E1+E5+E6+E7. *p < 0.05, **p < 0.01, ***p < 0.001.

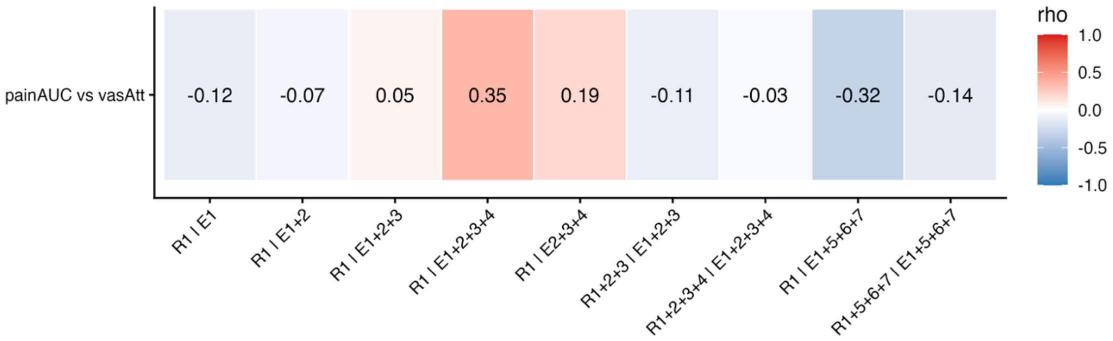


**Note:** Spearman coefficients ($r_s$) between pain AUC and attention ratings provided on Visual Analogue Scale. In general, the extent to which participants were able to focus on the target electrode (E1) during each of the trials were not related to pain AUC values – all presented coefficients were not significant.

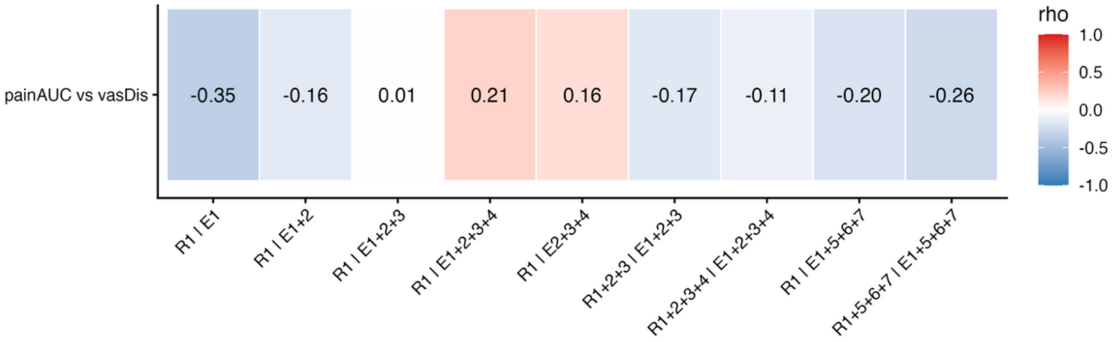


**Note:** Spearman coefficients ($r_s$) between pain AUC and extraction ratings provided on Visual Analogue Scale. In general, the extent to which they were able to extract pain from the target electrode (E1) versus the rest of electrodes during each of the trials were not related to pain AUC values – all presented coefficients were not significant.

**S5. Instructions on pain intensity, pain extraction and attentional focus on the pain experience.**

**i. Pain intensity:**

*"Please continuously rate the intensity of pain you experience throughout the entire period of stimulation. To provide your rating, use this device (trackball) by moving the ball to the left or right to draw a red line. The line should reflect the intensity of pain you feel in the specific location you are asked about. Please pay attention to the ends of the scale: on the left (no pain), on the right (the worst pain imaginable)."*

*"During the stimulation, adjust the length of the line so that it corresponds to the intensity of pain you are experiencing, using the reference points at both ends of the scale."*

**ii. Pain extraction:**

*"After each series, you will also be asked to answer two additional questions. You will provide your responses using the same device by drawing a blue line to the left or right and confirming your answer with the left button."*

*"The first question concerns how easy it was for you to extract the pain sensation from the specific location you were asked about during the stimulation from the overall pain sensation. Please pay attention to the ends of the scale: on the left (impossible), on the right (entirely possible)."*

**iii. Attentional focus on the pain experience:**

*"The second question concerns how easy it was for you to focus your attention on the pain sensation from the specific location you were asked about during the stimulation, compared to the overall pain sensation. Please pay attention to the ends of the scale: on the left (impossible), on the right (entirely possible)."*

**iv. Perceived locus of pain maximum:**

*"Was stimulation from any of the electrodes perceived as more painful than the others? If so, please indicate the electrode number."*

**S6. Quality checks: sensitivity under each electrode**

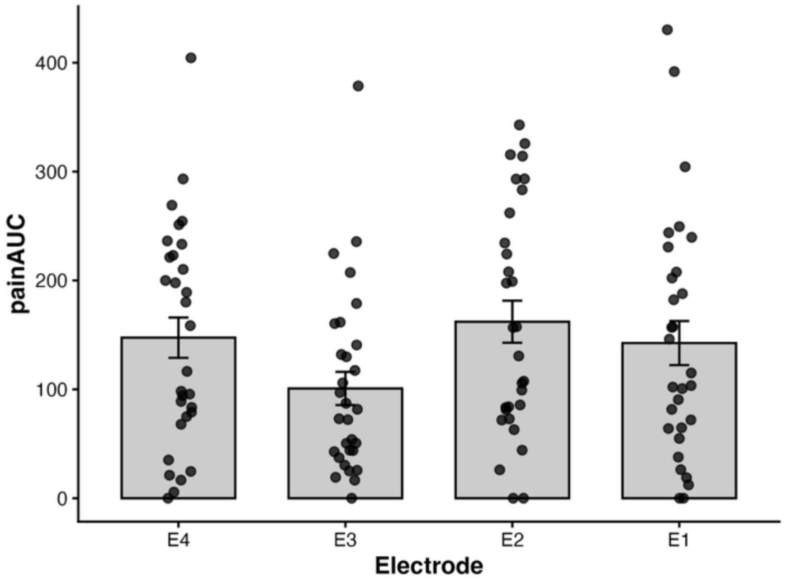


**Note:** Mean pain Area Under the Curve (AUC) values from block 1 of the paradigm in which only one electrode at the time was activated. Comparisons are presented in the Table below (S7).

**S7. Pain AUC (area under the curve) evoked by each distinct electrode (site)**

| Contrasts | Mean diff. | 95% CI | t | $p_{FDR}$ |
|---|---|---|---|---|
| E4 vs E3 | 46.67 | 07.74 to 85.60 | 2.45 | 0.06 |
| E4 vs E2 | -14.63 | -45.44 to 16.18 | -0.97 | 0.44 |
| E4 vs E1 | 4.98 | -30.80 to 40.76 | 0.28 | 0.78 |
| E3 vs E2 | -61.30 | -99.45 to -23.14 | -3.29 | 0.02 |
| E3 vs E1 | -41.69 | -84.64 to 1.27 | -1.98 | 0.11 |
| E2 vs E1 | 19.61 | -24.12 to 63.33 | 0.92 | 0.44 |

**Note:** E – electrode activated, FDR – false discovery rate, CI – confidence intervals

**S8. Exploratory robustness checks: FDR and Šidák corrected *p* values for all contrasts**

| Contrast | Trial 1 | Trial 2 | FDR | Šidák |
|---|---|---|---|---|
| H1.1 | LI0 | LI1 | 0.42 | 0.80 |
| H1.2 | LI0 | LI2 | 0.31 | 0.49 |
| H1.3 | LI0 | LI3 | < 0.001 | < 0.001 |
| H2.1 | LI0 | SHAM | 0.01 | 0.013 |
| H3.1 | LI1 | LI2 | 0.65 | 0.96 |
| H3.2 | LI2 | LI3 | < 0.01 | < 0.05 |
| H3.3 | LI1 | LI3 | < 0.01 | < 0.01 |
| H4.1 | LI0 | SSP3i | < 0.001 | < 0.001 |
| H4.2 | LI0 | SSP4i | < 0.001 | < 0.001 |
| H5.1 | SSP4i | SSP4b | 0.17 | 0.17 |
| E1 | LI3-SHAM* | LI0 | 0.03 | 0.13 |
| E2 | DNIC | LI3 | < 0.001 | < 0.001 |
| E3 | LI3 | SSP4i | < 0.001 | < 0.001 |
| E4 | DNIC | SSP4b | < 0.001 | < 0.001 |
| E5 | DNIC | LI0 | 0.77 | 1.00 |

**Note:** E – exploratory contrast, FDR – false discovery rate, CI – confidence intervals, H1 to H5 – hypotheses

**S9. Exploratory analyses to test for the effect of "block" and with covariate "sequence"**

| Model | Formula | Effect | *df* | *F* | *p* | $\eta_p^2$ |
|---|---|---|---|---|---|---|
| Original | DV ~ "trial" + (1 \| "subject") | "trial" | 8, 232 | 35.23 | *** | 0.55 |
| Additive | DV ~ "trial" + "block" + "sequence" + (1 \| "subject") | "trial" | 8, 500 | 53.36 | *** | 0.46 |
| | | "block" | 1, 500 | 9.60 | ** | 0.02 |
| | | "sequence" | 1, 500 | 2.70 | 0.10 | 0.01 |
| Interaction | DV ~ "trial" + "block" + "sequence" + "trial:block" + "trial:sequence" + (1 \| "subject") | "trial" | 8, 487.25 | 10.51 | *** | 0.15 |
| | | "block" | 1, 484.04 | 10.11 | ** | 0.02 |
| | | "sequence" | 1, 484.21 | 3.14 | 0.08 | 0.01 |
| | | "trial" × "block" | 8, 484.03 | 0.38 | 0.93 | 0.01 |
| | | "trial" × "sequence" | 8, 488.02 | 0.50 | 0.86 | 0.01 |

**Note:** The analyses were conducted to explore the potential role of habituation, under the assumption that the observed effects could be modulated by progressive habituation. We tested the effect of "block," which indicated that pain AUC values were overall significantly lower ($p < 0.01$) during the second repetition of the trials (block 4 vs. block 3). However, the effect of block did not interact with the "trial". Furthermore, the covariate "sequence" introduced in the interaction model also did not interact with the factor "trial".
